# FULL-LENGTH HYBRID TRANSCRIPTOME OF THE OLFACTORY ROSETTE IN SENEGALESE SOLE (*Solea senegalensis*): AN ESSENTIAL GENOMIC RESOURCE TO IMPROVE REPRODUCTION AT FARMS

**DOI:** 10.1101/2025.03.19.644159

**Authors:** Dorinda Torres, Andrés Blanco, Paula R Villamayor, Inmaculada Rasines, Ignacio Martín, Carmen Bouza, Diego Robledo, Paulino Martínez

## Abstract

Senegalese sole is a promising European aquaculture species whose main challenge is that captive-born males (F1) are unable to reproduce in farms, hindering breeding programs. Chemical communication through the olfactory system is hypothesized to stem this issue. Although significant advancement in genomic resources has been made over the past decade, scarce information exists on the genomic basis of olfaction, a special sensory system for demersal species like flatfish, which could play a prominent role in reproduction, social and environmental interactions. A full-length transcriptome of the olfactory rosettes including females, males, juveniles and adults, of both F1 and wild origins, was generated at the isoform-level by combining Oxford Nanopore long-read and Illumina short-read sequencing technologies. A total of 20,670 transcripts actively expressed were identified: 13,941 were known transcripts, 5,758 were novel transcripts from known genes, and 971 were novel genes encoding novel transcripts. Special attention was paid to the olfactory receptor gene families (OlfC, OR, ORA and TAAR) expression. Our comprehensive olfactory transcriptome of Senegalese sole provides a foundation for delving into the functional basis of this complex organ in teleost and flatfish. Furthermore, it provides a valuable resource for addressing reproductive management challenges in Senegalese sole aquaculture.

## 1. INTRODUCTION

Olfaction is a key sense for fish survival, as it plays a crucial role in fundamental behaviors such as reproduction, food detection, and interactions with other individuals and the surrounding environment in a broad sense [1,2,3,4]. Chemoreception through olfaction occurs within the olfactory organ, a paired structure known as the olfactory rosette, characterized by its multilamellar and rosette-shaped structure [5]. At cellular level, the olfactory epithelium comprises a complex arrangement of non-neuronal cells (supporting, basal and glandular cells) interspersed with olfactory sensory neurons (OSNs), including ciliated, microvillous, crypt, kappe and pear cell-types, the last two only described in zebrafish [6,7,8,9,10]. These OSNs express a single receptor gene through transcriptomic differences [11,12]. However, despite its well-known relevance as a fundamental sensory system, fish olfactory epithelium remains understudied, especially regarding intraspecific communication and its relevance for reproduction.

The olfactory receptors expressed in OSNs can detect a wide variety of semiochemical compounds, including pheromones. Although significant progress has been made in olfactory receptor deorphanization in the past decade [13,14,15,16,17], further research is still needed to elucidate the species-specific molecules that stimulate the different olfactory receptors, especially in non-model species [18,12]. OSNs generate chemical signals that are transmitted through their neuronal projections, ultimately converging to form the olfactory nerve, which reaches the brain for the first synapse at the level of the olfactory bulbs, the primary integrating centers of olfactory information [2,19].

Four main olfactory multigene families responsible of chemoperception have been described in teleost: olfactory receptors class C (OlfC), related to the mammal vomeronasal receptor V2R; odorant receptors (OR); olfactory receptors class A (ORA), related to the mammal vomeronasal receptor V1R; and trace amine-associated receptors (TAAR) [20,21,22,23,24]. Collectively, these families comprise the olfactory gene repertoire, with a wide variation in gene number among fish species [25]. Furthermore, a positive correlation has been shown between the structural complexity of the olfactory rosette and the size of the olfactory repertoire [26]. While important research has been conducted on the olfactory gene repertoire in model species such as zebrafish [20,21,27,28], scarce information is available for other teleost, especially for aquaculture species. Thus, understanding this complex sensory system is a first step for identifying potential pheromone-related socio-sexual behaviours. Altogether, this field of research provides valuable information for optimizing reproductive strategies and behavioral management in aquaculture [29].

Flatfish (Pleuronectiformes) constitute a diverse taxonomic group adapted to demersal lifestyle, including some species of great commercial value worldwide [30,31,32]. The development of genomic resources applicable to aquaculture breeding programs or sustainable fisheries management have greatly increased in the last decade [33,34,35]. The adaptation to a benthic environment, involving low light radiation and a sediment-rich seabed, made olfaction a critical and highly specialized sense for flatfish [36].

Senegalese sole (*Solea senegalensis*) is an emerging flatfish aquaculture species in Europe with great market value, which shows high growth and larval survival rates [37]. However, some bottlenecks, such as reproduction issues in captivity, still curtail the expansion of its production. Specifically, males born in farms (F1) present a behavioral dysfunction, being unable to court the females and carry out the subsequent fertilization of eggs [38,39,40,41]. Additionally, F1 males produce viable gametes but in lower volume than their wild counterparts [42,43]. Consequently, the production of *S*. *senegalensis* relies on the capture of wild males that are acclimated to captive conditions and utilized as breeders [44,39]. Thus, reproduction makes up one of the main constraints for developing breeding programs in this species, which has prompted diverse investigation approaches, including feeding [45,46], gamete production [47,43,48], testes methylation and transcriptome profiles [49], reproductive behavior [50,39,40,51,41], hormonal treatments [52,53,54,55], photoperiod rhythms [56,57,38], environmental enrichment [58], and chemical communication through olfaction [59,60]. While these studies contributed to a better understanding of the mechanisms controlling gonad maturation and reproduction in *S*. *senegalensis*, an explanation for the failure of courtship in F1 males has not been found yet. It has been hypothesized that chemical communication may underlie courtship failure and that environmental differences operating across life stages in farms *vs* wild may contribute to the low performance of F1 males [38,39,40]. Thus, studying the role of the olfactory system in courtship might provide crucial insights into the reproductive behavior issues of F1 males.

The arrival of new long-read sequencing technologies makes it feasible to obtain highly contiguous confident genomes [61,62], but also a much better characterization of transcriptomes at the isoform-level by combining short- and long-read sequencing using hybrid bioinformatic approaches [63,64,65]. The ability of the olfactory system to detect a huge variety of compounds suggests a complex underlying gene family repertoire, but also its resolution at the isoform level, especially in species with demersal lifestyle like flatfish [36].

The recent highly contiguous and annotated chromosome-level genome assembly of *S. senegalensis* [66,67] represents an essential resource for genomic exploration, including insights into genes associated with olfactory perception. However, the gene repertoire and transcriptome underlying the *S. senegalensis* olfactory organ remains scarce and, to our knowledge, only a transcriptomic profile of the olfactory organ comparing the dorsal rosette of males F1 *vs* males captured in the wild has been reported. Among the identified differentially expressed genes, some were either olfactory receptor genes or genes related to reproduction [59].

In the present study, a comprehensive and consistent isoform-level *S. senegalensis* olfactory transcriptome was generated by performing a hybrid sequencing approach, combining Illumina short-read and Oxford Nanopore long-read sequencing (ONT) technologies. Specifically, we conducted a transcriptomic comprehensive characterization of the olfactory rosettes of juvenile and adult individuals with different life-histories (female and male, F1 and wild origin), all acclimated to the same culture environment. We identified novel genes and vast transcript diversity, significantly improving the available genomic information of *S. senegalensis*. Our study represents the foundation for a confident investigation on the genetic mechanisms underlying the olfactory function in *S. senegalensis*, aiming to better understand its putative impact on the reproductive failure of F1 males, critical for application of breeding programs in this species.

## 2. MATERIALS & METHODS

### 2.1. Animal sampling and experimental design

A total of 15 animals were employed to characterize the transcriptome of the olfactory rosettes of *S. senegalensis*, six 10-months-old juveniles and nine 27 months-old adults. All fish were maintained indoors in tanks at the same standard water temperature and feeding conditions of the usual production protocol [68]. The main effort was put on long-read sequencing considering its higher resolution for reconstructing the full transcriptome, so 12 F1 individuals were used for this purpose: three juvenile males, three juvenile females, three adult males and three adult females. Additionally, we took advantage of a previous single-nuclei RNAseq dataset on the sole olfactory rosette, including three adult individuals acclimated to farm (F1 male, wild male and wild female) [69], to refine our olfactory transcriptome.

All animal procedures complied with the ethical regulations of the University of Santiago de Compostela responsible of the study, and the “Instituto Español de Oceanografía de Santander” and the company Stolt Sea Farm, which provided the animals for the study, in accordance with EU guidelines (86/609/EU). All fish were sacrificed by decapitation and immediately the two olfactory organs of each fish were dissected and preserved in liquid nitrogen, to be then stored at −80 °C until their use. Olfactory organs consist of a rosette-shaped paired structure located in the rostral part of the head, rostromedially to the eyes and close to the jaw. The olfactory rosettes are placed inside an olfactory chamber, enclosed by cartilage and coated by abundant connective tissue and mucous (Fig. 1). The dorsal and ventral olfactory chambers have direct communication with the environment through two nostrils that regulate the water-flow into the chamber. Both rosettes were dissected after accessing the olfactory chambers through an incision between the two nostrils.

**Fig. 1.**
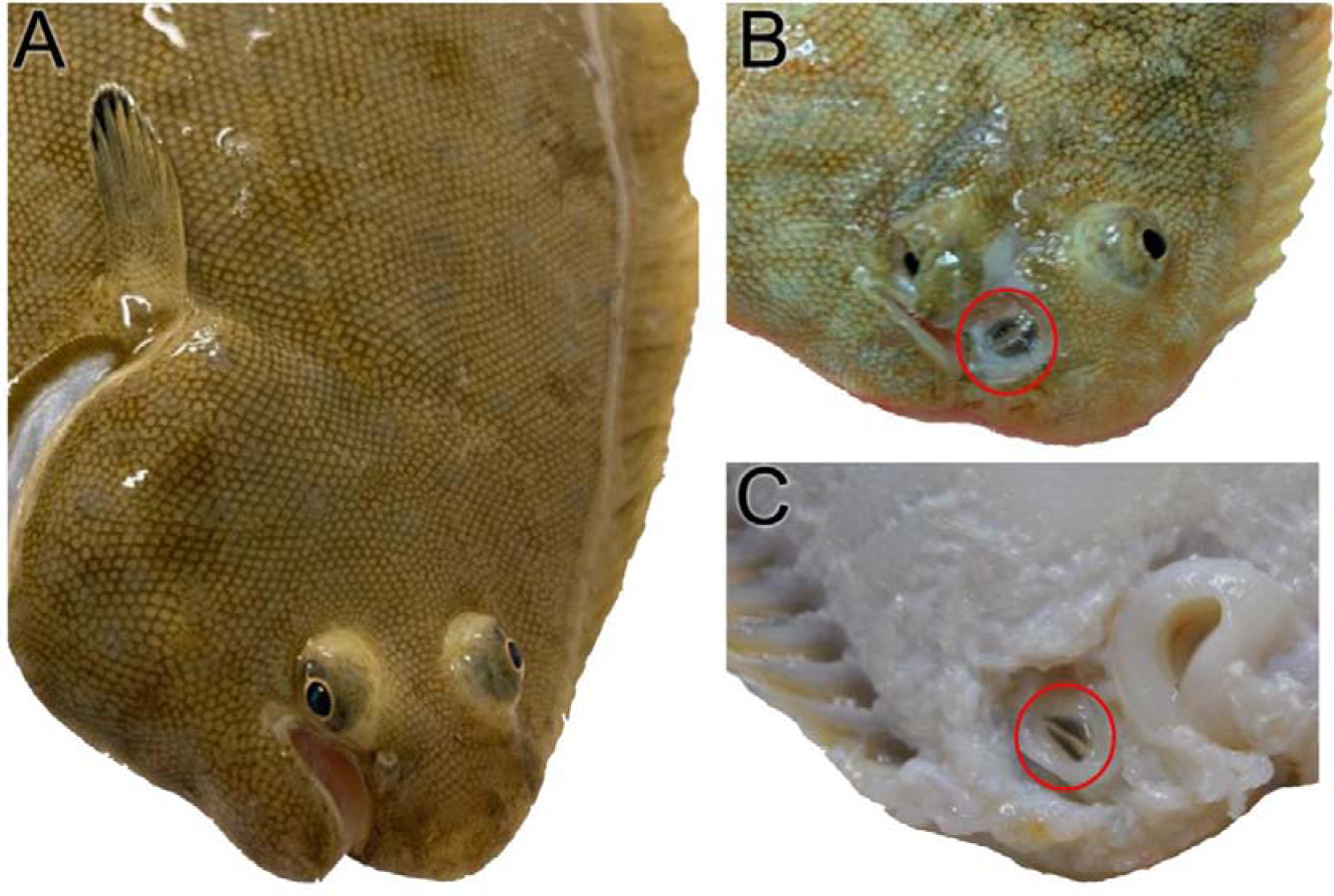
Macroscopic topography of the S. senegalensis olfactory rosettes. (A) Dorsal side, where both eyes are located, and the nostrils are visible close to the upper jaw. (B) Dorsal side after removing the skin that covers the olfactory chamber. Dorsal rosette (red circle) is strongly pigmented and surrounded by mucus. (C) Ventral blind side after removing the skin. Ventral rosette (red circle) with its black pigmentation.

### 2.2. Nanopore long-read sequencing

Total RNA extraction of dorsal and ventral rosettes for long-read sequencing was performed with TRIZOL Reagent (Life Technologies) according to manufacturer’s instructions. RNA quantity and quality were evaluated in a NanoDrop® ND-1000 spectrophotometer (NanoDrop® Technologies Inc.) and in a Bioanalyser 2100 (Agilent Technologies, “RNA integrity number” (RIN)). The 24 samples (12 dorsal olfactory rosettes and 12 ventral olfactory rosettes) were evenly pooled into a single sample (RIN = 9.6). The pool was delivered to Novogene (Cambridge, UK) for ONT library construction and long-read sequencing. The ONT raw reads were filtered with Nanofilt (https://github.com/wdecoster/nanofilt), and then aligned against the reference transcriptome of *S. senegalensis* (GCA_919967415.2) with minimap2 [70], applying the parameters established for ONT sequencing.

### 2.3. Illumina short-read sequencing

Olfactory rosettes’ Illumina sequences were obtained from nuclei RNA [69]. Briefly, nuclei were extracted [71], libraries were built in a 10X Chromium, and Illumina 150 bp paired end sequencing was performed in a NovaSeq 6000 at the Novogene Platform. The fasta files from raw sequencing were checked using FASTQC quality control. Reads from the six samples were aligned against the reference *S. senegalensis* genome (GCA_919967415.2) using STAR v2.7.9 [72], with default parameters.

### 2.4. Olfactory transcriptome characterization

For the olfactory transcriptome assembly, we combined ONT long-read and Illumina short-read sequencing datasets. *S. senegalensis* individuals from different origins (F1 and wild), sexes (males and females) and developmental stages (juveniles and adults) were gathered to achieve the broadest transcriptomic information as possible from diverse life-history specimens. This sequencing approach was expected to generate a comprehensive olfactory transcriptome at the isoform-level for the species.

Long-read data has become an essential tool for transcriptome reconstruction at the isoform level [73]. However, its integration with short-read data has demonstrated to be crucial for achieving a confident transcriptome reconstruction [74,65]. Therefore, the alignments used in our study from both sequencing techniques were merged by using the *--mix* option of StringTie v.2.2.1, since the hybrid dataset provides higher accuracy and sensitivity for gene and isoform identification [64]. The available genome annotation of *S. senegalensis* (GCA_919967415.2) was used as input. The resulting StringTie output comprised a GTF file containing information of the identified transcripts based on sequencing reads grouped by genomic coordinates. Transcripts were correlated, when possible, with known transcripts included in the annotation files of the *S. senegalensis* genome with their corresponding coding genes.

Several filtering steps were applied to the StringTie output to obtain a consistent dataset to be included in the *S. senegalensis* olfactory transcriptome. First, we performed a structural classification analysis of our transcripts by using SQANTI3 v.5.2.1 (https://github.com/ConesaLab/SQANTI3) [75], with default parameters. SQANTI3 provides a structural category and subcategory for each transcript based on how the reads match the available information in the annotated genome. The following categories were retrieved: i) Full splice match (FSM), an identical match to a known transcript in the genome annotation; ii) Incomplete splice match (ISM), all exons match to a known transcript, although some exons are missing at the ends; iii) Novel in catalog (NIC), a novel transcript with a new combination of known exons; iv) Novel not in catalog (NNC), novel transcripts containing at least one previously unknown exon; v) Intergenic, transcripts mapping to an intergenic region in the reference genome; vi) Genic, transcripts that overlap both known introns and exons in the reference genome; vii) Antisense, transcripts matching to the anti-sense strand of an annotated gene in the reference genome; and viii) Fusion, transcripts covering two different annotated genes.

Based on this categorization, we retained all FSM transcripts, while the rest of the transcripts were thoroughly evaluated before being included in the olfactory transcriptome of *S. senegalensis.* The filtering pipeline was applied to the entire StringTie output dataset to check that FSM transcripts were real transcripts that successfully passed the established filters.

First, we screened the StringTie v.2.2.1 transcript list obtained with RepeatMasker v.4.1.2 [76] using the bony fish (*Actinopterygii*) database, to identify putative repetitive elements (RE) in our transcripts. The identification of interspersed repeats representing transposable elements (TEs) within the coding sequences (CDS) was analyzed by cross-checking these data with the output from Transdecoder v.5.7.1 [77] utilizing default parameters. Transdecoder predicts CDSs by homology search against Pfam 33.0 to detect open reading frames (ORFs) with a minimum length of 100 amino acids. Subsequently, using an in-house Perl script, transcripts overlapping TEs more than 5% to the complete CDS were removed, but transcripts with trinucleotide simple repeats were kept in our dataset.

Once a consistent list of olfactory transcripts was obtained, we returned to the SQANTI3 structural annotation for in-depth analysis of the potential functionality of each transcript. SQANTI3 subcategories provide additional information for each transcript including exon number, coding region and presence of introns, among other data. Non-coding transcripts were removed. Furthermore, Antisense, Fusion and Genic categories (vi, vii and viii), were considered to not represent real olfactory transcripts, and thus, were discarded. Accordingly, only transcripts composed of mature RNAs expected to contain complete CDS and lacking introns were retained. Thus, the transcripts included within the subcategories “intron retention”, “mono-exon by intron retention” and “mono-exon” were removed. The latter was retained within the FSM category only when mono-exonic transcripts overlapped a mono-exonic known transcript. Finally, the mono-exon transcripts placed at intergenic regions were retained.

The FSM category was fundamental for *S. senegalensis* olfactory transcriptome reconstruction. The case where a transcript matched partially a reference isoform, was included within the ISM category. We filtered the ISM category by performing a pairwise comparison of our list with all FSM transcripts. When the transcript overlapped the FSM transcript of an annotated gene, we considered the ISM transcript of the pair pertaining to the FSM isoform. Consequently, ISM transcripts matching an existing FSM transcript of the same gene were not considered, since missing exons may be related to incomplete long-read sequencing. Another case within ISM category was when a transcript represented a novel isoform in which fewer exons than those described in the canonical FSM transcript (including all exons) were detected. In that case, these ISM transcripts were considered as novel and retained.

Moving into the categories composed of novel genes, the same strategy was followed for NIC and NNC, retaining novel transcripts that contained a novel combination of exons “combination_of_known_splicesites”, or one previously unknown exon “at_least_one_novel_splicesite”. While these subcategories were retained, the mono-exonic transcripts and those exhibiting introns were discarded. Intergenic transcripts, located in genomic regions where no information about coding regions existed in the reference genome were kept, representing novel genes.

### 2.5. Quantification of gene expression in the olfactory rosette

Normalized transcripts expression in the rosette was obtained as the number of transcripts per million of reads (TPM) provided by the StringTie output. There is lack of consensus regarding the TPM threshold, mostly depending on the sequencing technology employed, the specific tissue analyzed and the gene expression level [78,79,80]. We decided to use a TPM > 0.3 to consider a transcript as expressed in the olfactory rosette above the background noise considering the low expression levels of olfactory receptors [81,82,83,84] and that a high proportion of long-read sequences represented the whole transcript length (∼2kb the canonical transcript on average). This threshold could be similar to the 5 TPM usually used for short-read RNA-seq. The pipeline followed for the present study is shown in Fig. 2.

**Fig. 2.**
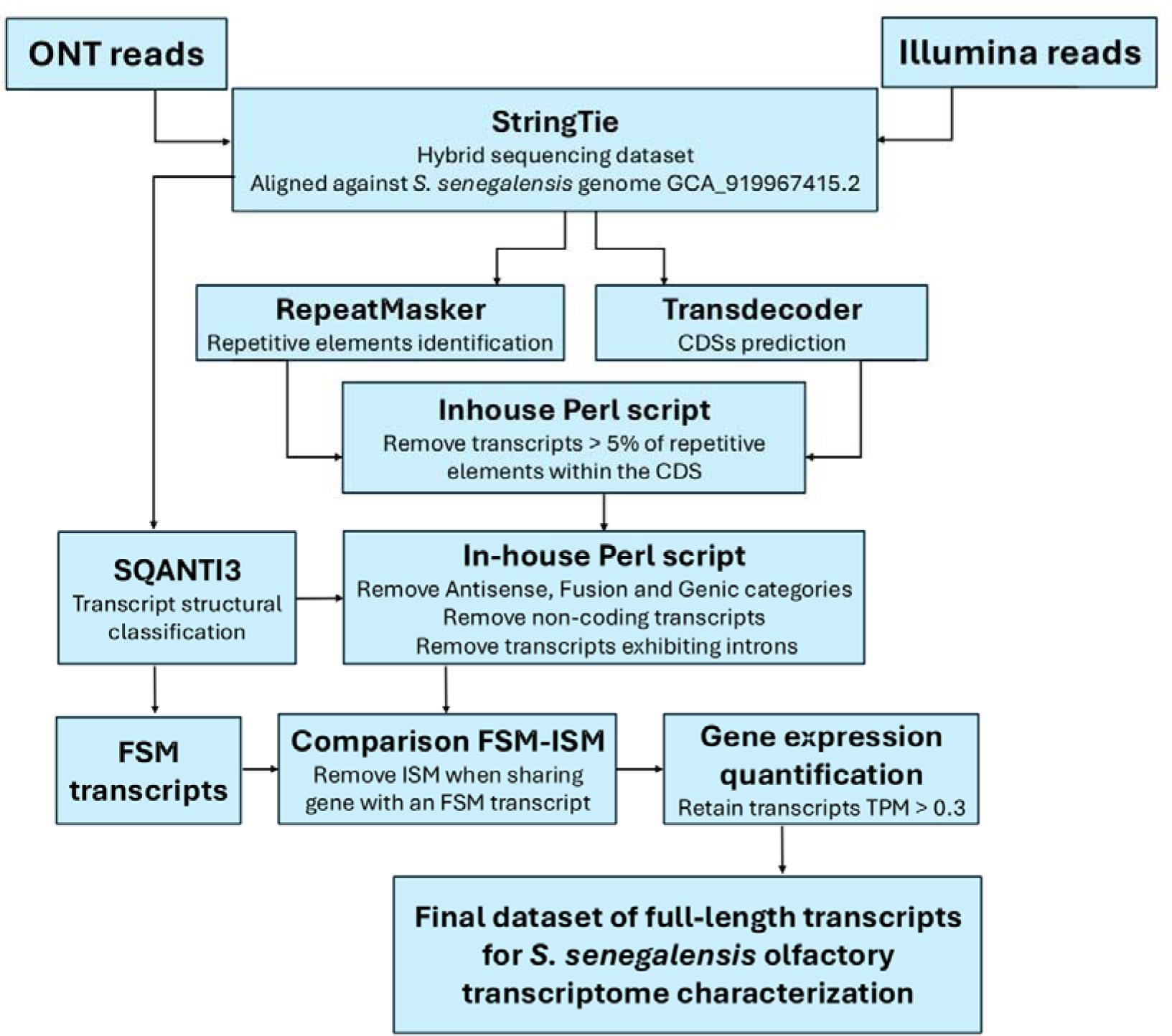
Overview of the pipeline for the olfactory transcriptome characterization. Full splice match (FSM); Incomplete splice match (ISM).

## 3. RESULTS & DISCUSSION

### 3.1. Olfactory transcriptome characterization

The chromosome-level genome assembly of *S. senegalensis* [66,67] has provided an invaluable resource for functional genomics evaluation of productive traits in this promising aquaculture species [37]. The olfactory rosette, responsible for chemical communication, has been recognized as an essential organ for courtship and reproduction [4]. However, to our knowledge, only a transcriptomic profile study explored its putative role on the reproduction failure of F1 males by comparing the transcriptome of olfactory rosettes of animals from wild and farm (F1) origin [59]. Additionally, in this study, a large differentially expressed gene repertoire was identified; as expected outcome considering the very different environmental conditions in the wild and farms. Indeed, to properly address the reproductive issue, the comparison of wild fish acclimated to farm environment with F1 fish would be necessary to identify the functional genomic signals underlying communication failure. The essential point for this analysis is to have available a high-quality reference olfactory transcriptome, especially considering the poor transcriptomic annotation of this complex sensorial organ in fish [36,85]. Such transcriptome was the primary goal of our study, and it was feasible through a long-read and short-read hybrid approach, essential to characterize the genes and isoforms involved in the olfactory function, particularly the olfactory receptor genes. Previous transcriptomic characterization through long-read sequencing techniques have been tackled in several tissues in flatfish species [86], but to our knowledge, this is the first olfactory transcriptome constructed through a hybrid sequencing approach in flatfish.

A total of 23.7 million (M) reads averaging 1,366 base pairs (bp) were generated through ONT long-read sequencing in our study from a single pool of dorsal and ventral olfactory rosettes including individuals of both sexes across various developmental stages. Of these, 98% mapped to the *S. senegalensis* reference genome. Illumina 150 bp pair-end reads from dorsal and ventral rosettes of three individuals with different life-histories ranged between 246 M and 540 M reads, 76.1% on average mapping to the reference genome (range: 71.8% to 82.3% reads per sample).

Our approach involved merging long- and short-read sequencing datasets to obtain a hybrid assembly using StringTie [74,64]. The inclusion of the Illumina short-read sequencing enriched the biological material for the analysis and enhanced the accuracy of exon and intron identification, reducing the errors inherent to ONT long-read sequencing, which allowed for more precise transcriptome characterization at the isoform level [65]. Although there are differences in third-generation sequencing techniques, both PacBio and ONT long-read sequencing are known to be suitable for transcriptomic analysis, and their combination with Illumina data significantly improves the results [87,65]. The StringTie output dataset rendered a total of 36,743 genes and 70,470 transcripts. Among them, 13,092 genes encoding 15,652 transcripts (22% of the total) were categorized as FSM by SQANTI3, representing a full match to the known transcriptome in the reference genome of the species [67].

All transcripts, including the FSM category, were further analyzed following an in-house pipeline based on previous results employing diverse sequencing techniques and bioinformatic tools in different taxonomic groups [88]. SQANTI3 software, as it provides accurate transcript structural annotation, greatly aided in the process of filtering transcripts from our dataset to obtain the final olfactory transcriptome [89,75]. Transdecoder, which identified the best candidate ORF for each transcript, aided to predict the complete CDS. Among the initial set of 36,743 genes, Transdecoder identified ORFs with more than 100 codons in 21,552 genes encoding 51,807 transcripts. Thus, these transcripts were predicted to contain a complete CDS, being capable of producing functional proteins.

RepeatMasker made it feasible to identify repetitive elements (RE) in the *S. senegalensis* transcriptome and to remove suspicious transcripts containing a significant portion of REs within the coding regions [76]. TEs have been demonstrated to be highly variable between species and to play a significant role in genome evolution [90], being involved in gene expression [91,92] or genome reorganization [93,94]. A remarkable variation in TE content has been reported in teleost, ranging from 5 to 56%, reflecting a positive correlation with genome size [95], which contrasts with the more constant RE proportion and genome size in other vertebrates such as mammals [96,97]. RepeatMasker detected 14.71% of REs in our transcripts, mainly involving retroelements and DNA transposons (12.57%) (Table 1). This value is similar to that reported in other fish transcriptomes [98], within the wide range reported across teleost (from 1.6% to 28% [99,100]). Further, the percentage of REs in the transcriptome was lower than that reported in the whole genome [101], an expected observation, considering the functional constraints of coding regions. Among TEs, the retroelements were represented in 5.68% of our transcripts, being LTR elements (3.39%) the most abundant, alongside LINEs (1.86%) and SINEs (0.43%). DNA transposons were present in 6.52% of our transcripts, being the hobo-Activator the most abundant (2.86%). Higher abundance of DNA transposons, or similar proportion of LINEs and DNA transposons was reported in other teleost species [102,100]. Other types of REs in our sequences with significant structural roles included simple repeats (1.51%), small RNAs (0.33%) and satellites (0.3%). We cross-checked the REs list retrieved from our sequencing data with the Transdecoder output to assess the overlapping of REs with ORFs in our transcripts, for an additional filtering. Transcripts were discarded from our olfactory transcriptome when REs constituted more than 5% of their CDS. At this point, our dataset consisted of a total of 36,703 transcripts encoded by 18,325 genes, averaging ∼2 transcripts per locus.

**Table 1.**
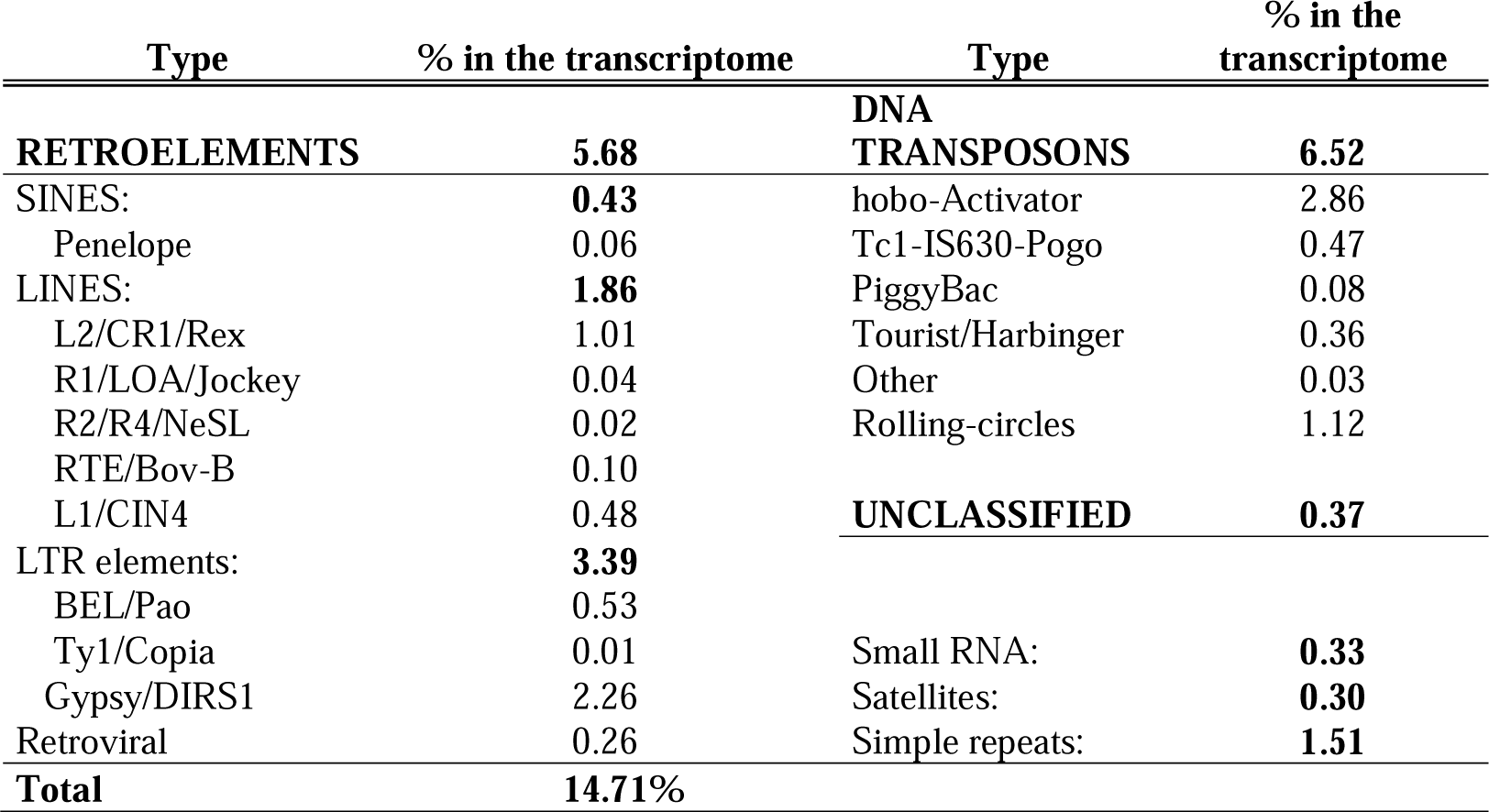
Percentage of repetitive elements overlapping the *S. senegalensis* olfactory transcripts using RepeatMasker.

Subsequently, we took advantage of the SQANTI3 categorization and discarded 2,342 transcripts included in the Antisense, Fusion and Genic categories to retain what we consider to be real transcripts matching our criteria. Also, we removed 1,568 transcripts classified as non-coding by SQANTI3 (Table 2).

**Table 2.**
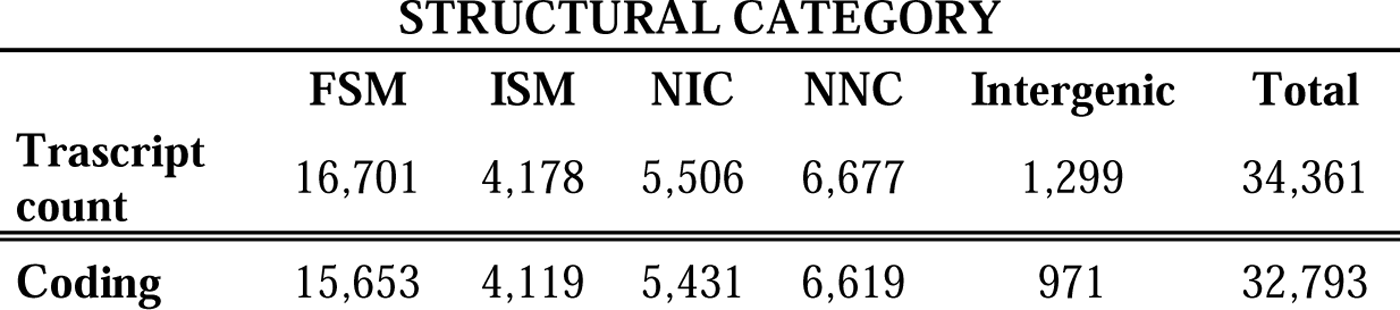
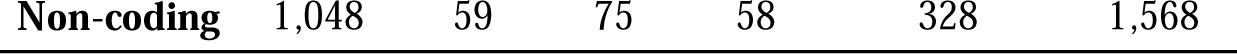
Structural categories in the olfactory transcriptome of *S. senegalensis*.

|  | STRUCTURAL CATEGORY |  |  |  |  |  |
| --- | --- | --- | --- | --- | --- | --- |
|  | FSM | ISM | NIC | NNC | Intergenic | Total |
| **Transcript count** | 16,701 | 4,178 | 5,506 | 6,677 | 1,299 | 34,361 |
| **Coding** | 15,653 | 4,119 | 5,431 | 6,619 | 971 | 32,793 |

Then, we explored the subcategories assigned to each of the 32,793 coding transcripts from which SQANTI3 provided valuable information of exon and intron distributions, and checked how these transcripts matched the available information in the reference genome (Table 3). Transcripts that matched to a mono-exonic transcript in the genome were included in the “mono-exonic” subcategory of FSM and thus, retained. Conversely, if a mono-exonic transcript matched a reference multiexonic transcript, it was discarded due to missing information, except for mono-exonic transcripts within the intergenic category, for which no previous information existed.

**Table 3.**
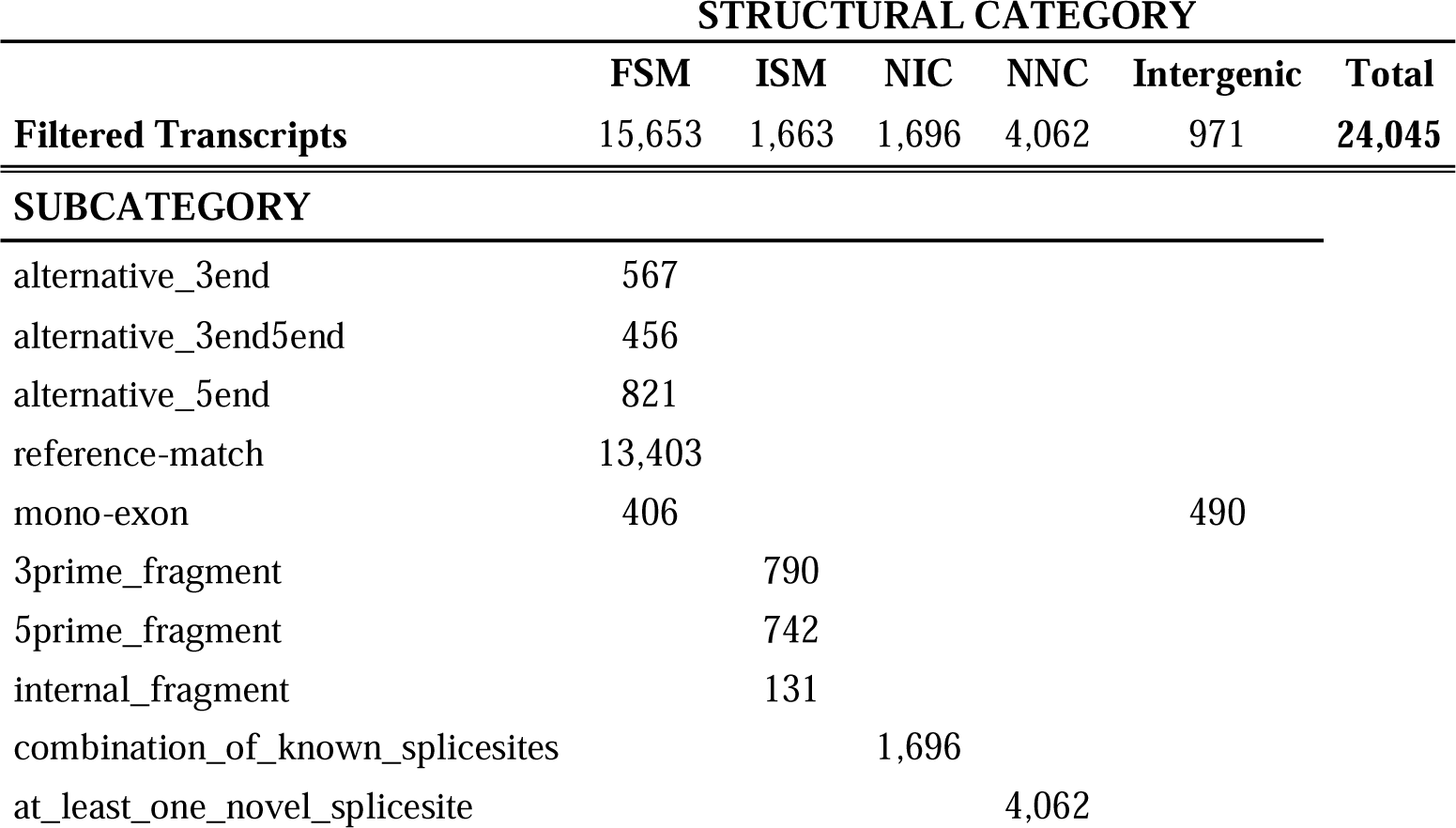
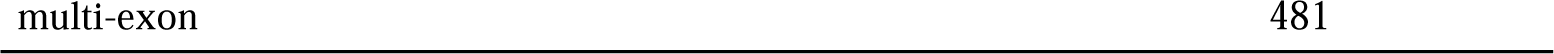
Structural transcript categories and subcategories for olfactory transcripts of *S*. *senegalensis*.

|  | STRUCTURAL CATEGORY |  |  |  |  |  |
| --- | --- | --- | --- | --- | --- | --- |
|  | FSM | ISM | NIC | NNC | Intergenic | Total |
| **Filtered Transcripts** | 15,653 | 1,663 | 1,696 | 4,062 | 971 | **24,045** |
| **SUBCATEGORY** |  |  |  |  |  |  |
| alternative_3end | 567 |  |  |  |  |  |
| alternative_3end5end | 456 |  |  |  |  |  |
| alternative_5end | 821 |  |  |  |  |  |
| reference-match | 13,403 |  |  |  |  |  |
| mono-exon | 406 |  |  |  | 490 |  |
| 3prime_fragment |  | 790 |  |  |  |  |
| 5prime_fragment |  | 742 |  |  |  |  |
| internal_fragment |  | 131 |  |  |  |  |
| combination_of_known_splicesites |  |  | 1,696 |  |  |  |
| at_least_one_novel_splicesite |  |  |  | 4,062 |  |  |

The maturation of mRNA involves several processes, including splicing, where non-coding introns are removed from primary mRNAs capable of translating into a protein. However, intron retention has sometimes been demonstrated to influence the regulation of gene expression [103,104], and a certain proportion of immature mRNAs including introns may be identified [105]. We found that 22% of our transcripts still included introns. We cannot conclude whether these transcripts containing introns play a significant role in the regulation of gene expression or just represent immature mRNAs [106]. Thus, we removed 7,421 transcripts subcategorized as “intron retention” and “mono-exon by intron retention”.

At this point of the filtering process, our *S. senegalensis* olfactory transcriptome consisted of five structural categories according to their correspondence with the available genome information. FSM, representing a perfect match to a known transcript, included 85.60% transcripts classified as “reference match”. The rest of FSM transcripts included slight variations, either missing 3’ end or 5’ end, or both. The explanation behind the variability between transcript boundaries may be associated with the poor annotation of the UTR regions in the species. In the same way, ISM transcripts were associated by SQANTI3 with a known transcript in the genome, although some exon(s) were missing compared to the reference isoform. Special attention was paid to this category, since ISM transcripts could represent novel transcripts with fewer exons than the canonical FSM. The pairwise comparison between FSM and ISM revealed a total of 1,191 pairs sharing the same associated gene and transcript. Following a conservative criterion, we assumed that those ISM corresponded to the same FSM transcript. Still, we retained the remaining 1,663 ISM transcripts that were associated with a single gene lacking an FSM transcript in our dataset. These transcripts need to be further explored to know whether they represent novel transcripts with a novel exon combination, or whether they are artifacts of ONT sequencing.

For the examination of categories NIC and NNC composed of novel transcripts not annotated in the reference genome, we considered the scarce and puzzling annotation of the olfactory genes in teleost, and specifically in *S. senegalensis*. Both categories consisted of known genes with a novel isoform, either by a new combination of known splice-sites or by the appearance of a new splice-site. The NIC category included 1,700 transcripts consisting of new combinations of annotated exons, representing novel transcripts of a known gene. We found a larger number within the NNC category, where 4,103 novel transcripts that contained at least one previously unannotated exon were identified. These findings highlight the potential of long-read RNA sequencing for identifying novel transcripts, expanding our understanding of transcriptome complexity[107].

The Intergenic category included novel transcripts placed in a genomic region with no previous annotated genes, which was possible by combining long-read with Illumina sequencing data using a broad sampling collection across different life-history fish. At this point, the *S*. *senegalensis* olfactory transcriptome consisted of 24,045 transcripts distributed in different categories and subcategories (Table 3).

### 3.2. Gene expression in the olfactory epithelium

Given the conservative filtering pipeline followed in our study, it can be assumed that we retained real transcripts with an active function in the olfactory rosette of *S*. *senegalensis*. To estimate if the transcripts identified were expressed in the olfactory rosette, a threshold of TPM > 0.3 from StringTie TPM values was set (see Materials & Methods section).

A total of 20,670 full-length transcripts encoded by 14,917 genes (1.39 isoforms per gene) were expressed in the *S. senegalensis* olfactory transcriptome according to this criterion (Table 4); 14,708 were protein coding genes and 209 were classified as long non-coding RNA (lncRNA) genes. Long-read sequencing has become the preferred approach for comprehensive characterization of lncRNAs [108], a gene class that can modulate chromatin function, the assembly and function of nuclear bodies, the stability and translation of cytoplasmic mRNAs and that can interfere with signaling pathways [109]. A more detailed description for each transcript included in our olfactory transcriptome is given in Supplementary Table 1.

**Table 4.**
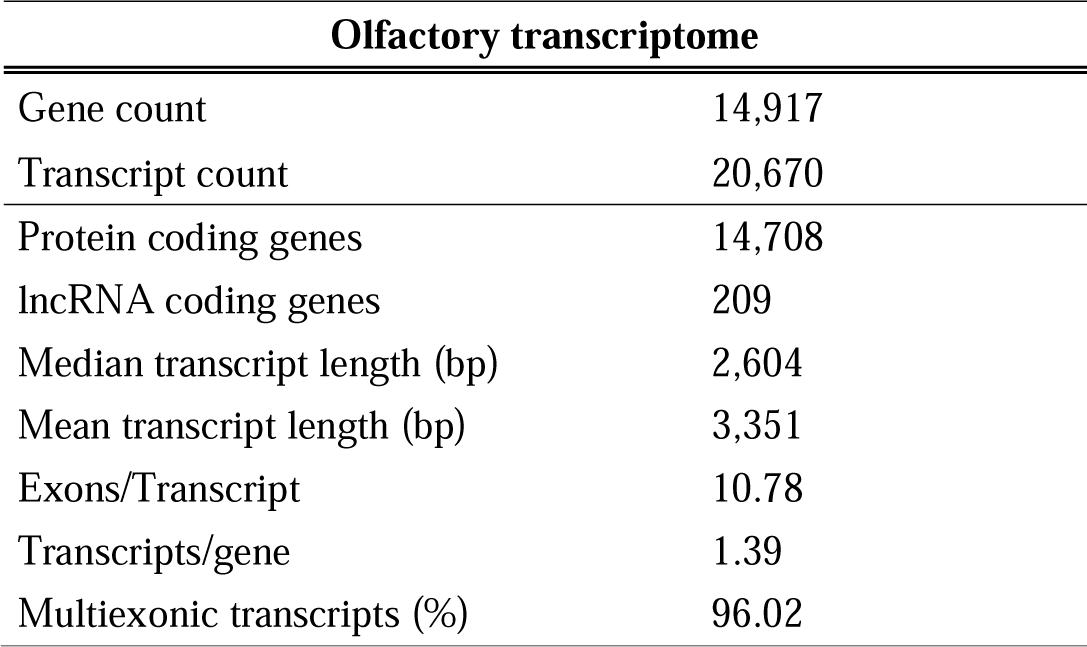
Summary statistics of the S. senegalensis olfactory transcriptome.

| Olfactory transcriptome |  |
| --- | --- |
| Gene count | 14,917 |
| Transcript count | 20,670 |
| Protein coding genes | 14,708 |
| lncRNA coding genes | 209 |
| Median transcript length (bp) | 2,604 |
| Mean transcript length (bp) | 3,351 |
| Exons/Transcript | 10.78 |
| Transcripts/gene | 1.39 |
| Multiexonic transcripts (%) | 96.02 |

From the 14,917 genes constituting the olfactory transcriptome, 11,925 were annotated in the current genome of the species, whereas 2,021 possessed an Ensembl ID but no annotation. The remaining 971 were novel genes identified through our transcriptome characterization, and therefore, lacked Ensembl ID and annotation.

*S*. *senegalensis* genome possessed 190 genes annotated as olfactory receptors, comprising 72 OlfC, 48 OR, 3 ORA, and 67 TAAR. Among them, 143 were expressed in the olfactory rosette of the individuals under studied, and therefore, included in our olfactory transcriptome; distributed as follows: 64 OlfC, 35 OR, 2 ORA, and 42 TAAR. These genes encoded 191 transcripts actively expressed (TPM > 0.3) in the olfactory rosette of the individuals studied. These receptors that comprise the olfactory repertoire are responsible for odor detection, and therefore, play a crucial role in chemical communication through olfaction [4,12]. Although significant progress has been made in identifying molecules that stimulate specific olfactory receptors in various species [13,14,15,16,17], no deorphanization studies have been conducted for *S. senegalensis* olfactory receptors. Hence, future studies should focus on identifying potential chemical cues that stimulate olfactory receptors in this species to optimize reproductive strategies in aquaculture [29].

All in all, we retained 12,278 FSM transcripts, 1,663 ISM transcripts, 1,696 NIC transcripts, 4,062 NNC transcripts and 971 intergenic transcripts in the *S. senegalensis* olfactory transcriptome (Figs. 3A and 3B), 67.4% of them constituting known transcripts (FSM and ISM categories) included in the current genome of the species. The rest were novel transcripts (NIC, NNC and Intergenic categories) that significantly improved the annotation of the species genome.

**Fig. 3.**
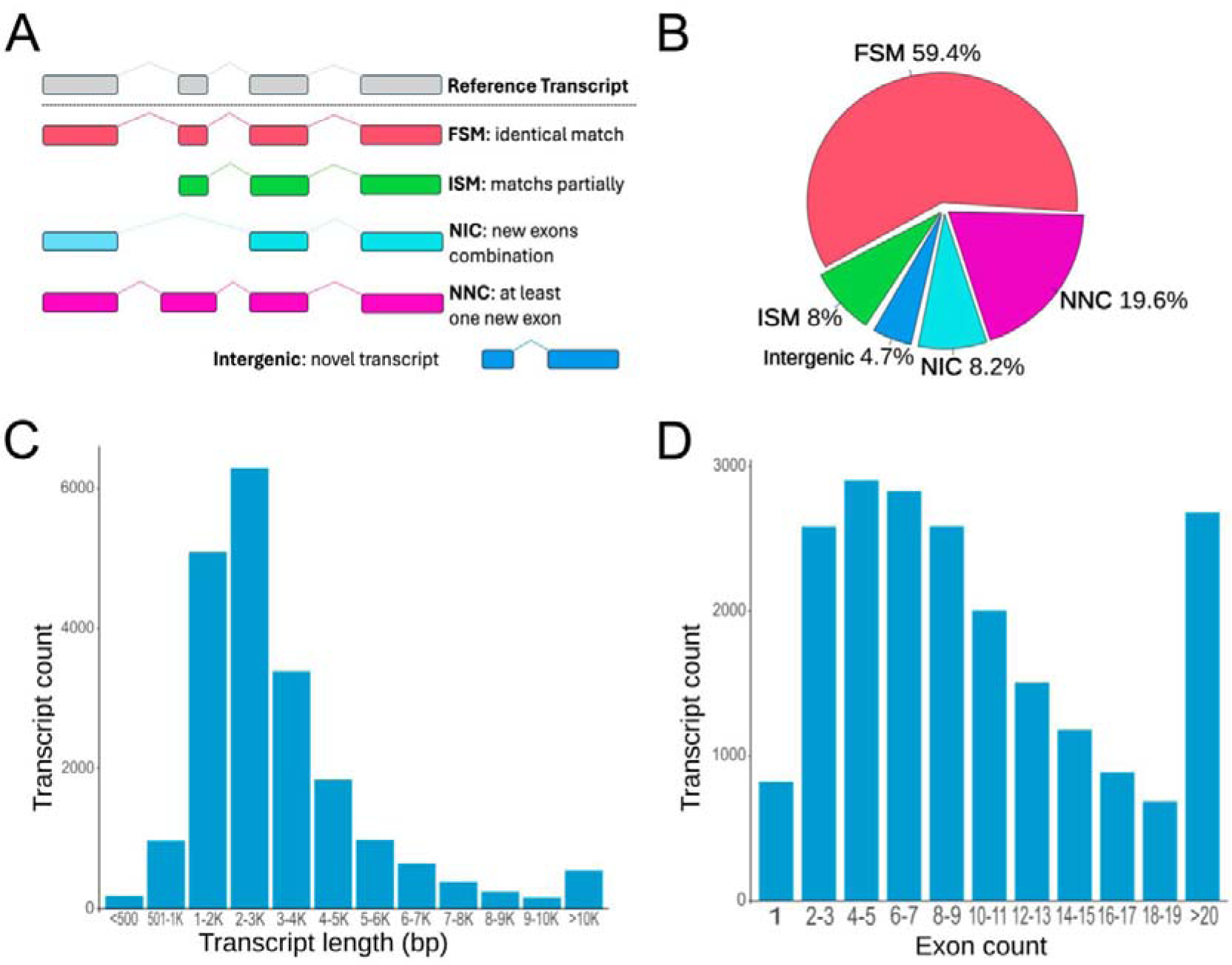
Main features of the olfactory transcriptome of S. senegalensis: (A) Structural categories based on the reference transcript genome information; (B) Distribution of structural categories: full-splice match (FSM), incomplete splice match (ISM), novel not in catalog (NNC), novel in catalog (NIC) and intergenic; (C) and (D) Transcript length and Exon number distribution, respectively.

Distribution transcript length ranged from 303 to 90,883 bp, with a median of 2,604 bp and a mean of 3,351 bp, 38 transcripts being above 30 kb (Fig. 3C). Gene size ranged between 249 and 666,329 bp, with a median of 8,922 bp and a mean of 18,212 bp, 328 genes being above 100 kb. This represents a larger median gene length than the 7,566 bp reported for the whole genome [67]. Gene and transcript length are positively correlated, with a trend of increased length in highly conserved genes [110,111]. Furthermore, significant variations in gene length have been observed across tissues, with longer transcripts predominantly expressed in blood vessels, nerves and brain, while shorter transcripts are more commonly found in skin and gonads [112]. The olfactory rosette is a complex highly vascularized organ with a significant composition of neural cells [7], and therefore, it would be expected to express longer transcripts. Most transcripts (96%) were multiexonic, exhibiting a mean of 10.8 exons and a median of 8.0 exons per transcript. A total of 2,682 transcripts consisted of ≥ 20 exons, while 4-5 exons were the modal class in our dataset, including 2,902 transcripts (Fig. 3D).

Expressed genes within the olfactory transcriptome, besides a unique StringTie code assigned, were associated, when possible, to an annotated gene in the reference genome. This rendered 13,946 known genes in the *S*. *senegalensis* olfactory transcriptome and the discovery of 971 novel genes, significantly enriching the current number of protein coding genes in Ensembl rapid release. Among the total olfactory transcript count, 13,941 (67.4%) were known full-length annotated transcripts in the genome, while 6,729 were identified as novel transcripts in our study. Among these, 5,758 transcripts (27.9%) were alternative splice variants encoded by known genes, whereas 971 (4.7%) were encoded by novel genes located on intergenic regions, not registered in the reference genome.

We identified a high proportion of known genes and transcripts actively expressed in the olfactory rosette, and furthermore, we significantly enriched gene and transcript count of the species, adding to the list relevant genomic information related to olfaction. Thus, despite the olfactory transcriptome should still be enriched by exploring new developmental stages and environmental conditions, this information represents a foundation for investigating the role of the olfaction in social communication and reproduction, where the full-length mRNA sequencing proved its potential to uncover an unprecedent number of novel transcripts emerging from alternative transcription initiation or splicing [113,79]. The substantial number of newly identified transcripts is consistent with findings reported in analogous studies [114,88]. Our study presents a confident and comprehensive olfactory transcriptome at the isoform level in *S. senegalensis*. It represents the first hybrid sequencing approach for olfactory transcriptome characterization in any flatfish species, a critical organ for demersal species. This approach enabled the identification of novel genes and transcripts, significantly enriching the genomic information in the species. The *S. senegalensis* olfactory transcriptome represents a foundation for understanding how this complex sensory system operates in teleost species. Future research will benefit from integrating our transcriptomic information with functional studies that might help to solve the reproductive dysfunction in the species contributing to improving fish aquaculture production.

## ACKNOWLEDGMENTS

We acknowledge the technical support and informatic resources provided by the Centro de Supercomputación de Galicia (CESGA). This study was funded by: PID2022-137821OB-C31 funded by MICIU/AEI/10.13039/501100011033 and by “ERDF/EU”; RSE Saltire International Collaboration Award (1856); Consellería de Economía, Industria e Innovación e Consellería de Cultura, Educación, Formación Profesional e Universidades, Xunta de Galicia (06_IN606D_2022_2693134; ED431C 2022/33); Proxectos Colaborativos Campus Terra-USC (SOLREP/2022-PU015).

## Ethics approval

All animal experiments were conducted in accordance with the guidelines of the University of Santiago de Compostela, the Instituto Español de Oceanografía de Santander and Stolt Sea Farm, in accordance with EU guidelines (86/609/EU).

## Author contributions

DT: Conceptualization, Investigation, Data curation, Formal analysis, Writing – original draft; ABH: Software, Formal analysis; PRV: Conceptualization, Investigation, Formal analysis, review and editing; IM&IR: Resources, review and editing; CB: review and editing; DR: Conceptualization, Funding acquisition, review and editing; PM: Conceptualization, Funding acquisition, Project administration, Supervision, review and editing Methodology, Writing – original draft, review and editing.

## Data availability

The datasets generated during the current study will be made available upon acceptance of the manuscript.

